## supplemental java program 2 for "Comparing Theories for the Maintenance of Late LTP and Long-Term Memory: Computational Analysis of the Roles of Kinase Feedback Pathways and Synaptic Reactivation"

```
import java.io.*;
import java.util.Random;

public class fig3b

 {  public static void main (String args[]) throws IOException {

    PrintWriter out1 = new PrintWriter(new FileWriter("ck2act.txt"));
    PrintWriter out2 = new PrintWriter(new FileWriter("erkact.txt"));
    PrintWriter out3 = new PrintWriter(new FileWriter("pkaact.txt"));
    PrintWriter out4 = new PrintWriter(new FileWriter("wsyn.txt"));
    PrintWriter out5 = new PrintWriter(new FileWriter("tagsyn.txt"));
    PrintWriter out6 = new PrintWriter(new FileWriter("prew.txt"));
    PrintWriter out7 = new PrintWriter(new FileWriter("tagp1.txt"));
    PrintWriter out8 = new PrintWriter(new FileWriter("tagp2.txt"));
    PrintWriter out9 = new PrintWriter(new FileWriter("tagp3.txt"));
    PrintWriter out10 = new PrintWriter(new FileWriter("casyn.txt"));
    PrintWriter out11 = new PrintWriter(new FileWriter("rafp.txt"));
    PrintWriter out12 = new PrintWriter(new FileWriter("pck2.txt"));
    PrintWriter out13 = new PrintWriter(new FileWriter("perk.txt"));
    PrintWriter out14 = new PrintWriter(new FileWriter("pkms.txt"));

    double dt=0.000025; // high-res timestep (min)
    double delta=0.01; // low-res timestep (min)

double recstart=1000.0; // Time to start writing data

    double stime=recstart+2500.0; // Time of stimulus onset

    double isi=5.0;      // Spacing between tetanic stimuli

    double recend=stime+3000.0;

    int recintvl=20; // (delta) Intervals to record

    double tref; // Can be used multiple places in program, for example to denote time since inhibitor onset, as long as it is reset as needed.

	double mnreact=500.0;
	double stdreact=150.0;
	double reactint=mnreact;
	int kreact=0;

//PROGRAM VARIABLES

// Activities of kinases other than MAPK cascades
	double ck2act;
	double camp;
	double pkaact;

// MAPK CASCADE
	double raf;
	double rafp;
	double mkk;
	double mkkp;
	double mkkpp;
	double erk;
	double erkp;
	double erkpp;
	double erkact;

// SYNAPTIC VARIABLES
	double tagp1;
	double tagp2;
	double tagp3;
	double tagsyn;
	double prew;
	double wsyn;
	double wbas=0.0;

// SIMPLE PLASTICITY RELATED PROTEIN
	double prp;

// pkm level, and phosphorylation sites on TFs for CaMKII and ERK

	double pck2;
	double perk;
	double pkms;

// TIME DERIVATIVES

    	double dv1dt;
    	double dv2dt;
    	double dv3dt;
    	double dv4dt;
    	double dv5dt;
    	double dv6dt;
    	double dv7dt;
    	double dv8dt;
    	double dv9dt;
    	double dv10dt;
    	double dv11dt;
    	double dv12dt;
    	double dv13dt;
    	double dv14dt;
	double dv15dt;
	double dv16dt;
	double dv17dt;
	double dv18dt;
	double dv19dt;
	double dv20dt;

// Cytoplasmic and nuclear Ca concentrations, camp function
	double casyn;
	double powca;
	double powkc;
	double termcamp;

// WORK IN CONCENTRATION UNITS OF uM, TIME UNITS OF MIN

// MODEL PARAMETERS FOLLOW:

// BASAL PKA AND CA, STIMULUS AMPLITUDES. CA2 refers to nuclear Ca.
// Stim amps also drive PKA elevations.

	double Cabas=0.04;
	double campbas=0.06;
	double AMPTETCA=0.8;
	double cadur=0.05;

	double AMPTETCMP=0.25;
	double dcamp=1.0; // duration of tetanic cAMP elevation. 1.0 min standard RTS, rolipram cures RTS-basal by raising dcamp to 2.0.

	double AMPTETSTIM=0.13;

	double AMPTETREACT=0.08; // maximal Raf reactivation stim amplitude.  Below this multiplies with coupling parameter, epsilreact = 0.3, to give the 0.024 value in the paper for reactivation Raf amplitude.

	double rasdur=1.0; // 1.0 standard

	double ampstim; // set later to Raf activation stimulus amplitude

// RATE CONSTANTS AND MICH CONSTANT FOR CAMKIV AND CAMKII ACTIVATION
	double kfck2=180.0; // 180 standard for most sims

	double tauck2=1.0;
	double Kck2=0.7;

//CONSERVED ENZYME AMOUNTS
	double raftot=0.25;
	double mkktot=0.25;
	double erktot=0.25;

// MAPK rate and Michaelis constants
	double kbraf=0.12;
	double kfmkk=0.6;
	double kbmkk=0.025;
	double Kmkk=0.25;
	double kferk=0.52;
	double kberk=0.025;
	double Kmk=0.25;
	double kfbasraf=0.0075; /* 0.0075 standard for most sims */

// Inhibition factors for enzymes and for plasticity related protein gene

	double inherk;
	double inhmkk;
	double inhck2;
	double inhgp;
	double inhpka;
	double inhpkm;

// Rate constants and other parameters
// describing PKA activation and synaptic tagging

	double Kcamp=1.0;

	double kphos1=0.15;
	double kdeph1=0.008;
	double kphos2=0.8;
	double kdeph2=0.2;
	double kphos3=0.06;
	double kdeph3=0.05;
	double taupka=15.0;

// PKM parameters

	double kphos4=0.1;
	double kdeph4=0.1;
	double kphos5=2.0;
	double kdeph5=0.1;
	double kdegpkm=0.02;

	double ktranspkm1=0.2;

	double Kpkm=0.75;
	double vbaspkm=0.0015;

// Maximal and basal transcription rates for transcription of plasticity related protein (could also be interpreted as translation), and for decay of PRP.

// Note that inhibition factor for protein synthesis or transcription is given a value here

	double vsyngp=0.01;

	inhgp=1.0;

	double taugp=100.0;

// Rate constants and other parameters for synaptic weight changes

/* kltp=500, Kpr2=0.2, vprew=0.0035, for single decaying tetanic LTP, no feedback. */


/* kltp = 500 for generating bistability in W with the upper state stabilized by synaptic reactivation. */

	double kltp=480.0;

	double vsynbas = 0.00086;

	double tausyn=3200.0; // ten times longer for this model variant.

	double kprew=6.0;

	double Kpr2=0.2; // Dissociation type constant by which a decrease in PREW protein suppresses rate of increase of W.

	double vprew=0.0035;

	double tauprew=100.0;

// Parameters for coupling Ca transients to synaptic weight in reactivation plus feedback

	double Ksyn=4.0;

	double epsilreact=1.0; /* Essential coupling constant between reactivation of synapse and bistability.  For simulating molecular feedback loops only, set to 0.0 */

	double test1;
	double test2;
	double rint1;
	double rint2;
	double rndbm;
	double synhill;


// TIME STUFF, COUNTERS, VALUE ARRAY

    double time=0.0; // time (minutes)

	double treact=0.0;  // time variable reset at each reactivation

    int i, k, j, l, m, iout, ibig; // counters

    double[] values=new double[20];


// VARIABLE INITIALIZATION. Use nonzero initial values
// to avoid extremely small numbers in output files.

/* for slower variables, initial conditions are set reasonably close to equilibrium basal values, for fast variables 0.001 is fine */

	values[1]=0.001;
	values[2]=0.001;
	values[3]=0.5*raftot;
	values[4]=0.3*mkktot;
	values[5]=0.4*mkktot;
	values[6]=0.3*erktot;
	values[7]=0.4*erktot;
	values[8]= 0.001;
	values[9]= 0.001;
	values[10]= 0.001;
	values[11]=inhgp*vsyngp*taugp;
	values[12]=vprew*tauprew;
	values[13]=vsynbas*tausyn;
	values[14]= 0.001;
	values[15]= 0.001;
	values[16]= 0.001;

// MAIN LOOP (LARGER TIMESTEP, IF TIME COURSE DATA IS OUTPUTTED THIS LARGER TIMESTEP IS USED)

        k=1;
        do {

// INNER SIMULATION LOOP (SMALLER TIMESTEP DT)

            j=1;
            do {

		tref=time-stime;
		casyn=Cabas;
		ampstim=0.0;
		camp=campbas;

		ck2act=values[1];
		pkaact=values[2];

		raf=values[3];
		mkk=values[4];
		mkkpp=values[5];
		erk=values[6];
		erkpp=values[7];

		tagp1=values[8];
		tagp2=values[9];
		tagp3=values[10];

		prp=values[11];

		prew=values[12];
		wsyn=values[13];

		pck2=values[14];
		perk=values[15];
		pkms=values[16];

// IMPOSE A TRAIN OF CA TRANSIENTS, AND ELEVATED CAMP AND KFRAF

/* The nonlinearity of the reactivation feedback is mostly because the amplitude of the calcium and camp transients during reactivation depend on this function, and reinforcement of synaptic strength is very sensitive to these amplitudes */

		casyn=Cabas;
		ampstim=0.0;
		camp=campbas;
		synhill=(wsyn*wsyn*wsyn*wsyn*wsyn/(wsyn*wsyn*wsyn*wsyn*wsyn + Ksyn*Ksyn*Ksyn*Ksyn*Ksyn));


	if (treact > reactint)
       {
       treact=0.0;
       kreact=1;
       }

	if ((treact < cadur) && (kreact==1))
		{
		  casyn=epsilreact*synhill*AMPTETCA;
		  if (Cabas > casyn) {casyn=Cabas;}
		}

	if ((treact < dcamp) && (kreact==1))
		{
		  camp=epsilreact*synhill*AMPTETCMP;
		  if (campbas > camp) {camp=campbas;}
		}

	if ((treact < rasdur) && (kreact==1))
		{
		  ampstim=epsilreact*synhill*AMPTETSTIM;
		}

	if ((treact > rasdur) && (kreact==1))

		{
		  kreact=0;

/* use Box-Mueller algorithm to generate a Gaussian random variable for the time between reactivations, reactint, from two uniformly distributed (on 0,1) random variables */
		  test1=Math.random();
		  test2=Math.random();
		  rint1=-2.0*(Math.log(test1));
		  rint2=Math.sqrt(rint1);
		  rndbm=rint2*(Math.cos(2.0*3.141592654*test2));
		  reactint=mnreact+stdreact*rndbm;
		  if (reactint < 10.0) {reactint=10.0;}
		}


// SIMPLE SQUARE WAVES FOR CYTOPLASMIC CA

/* note that several LTP stimulus protocols, including chem-LTP and theta-LTP, can be simulated here. */

// Tetanic protocol

		if ((time-stime) > 0.0 && (time-stime) < cadur)
		  {
		  casyn=AMPTETCA;
		  }
		if ((time-stime) > 0.0 && (time-stime) < dcamp)
		  {
		  camp=AMPTETCMP;
		  }
		if ((time-stime) > 0.0 && (time-stime) < rasdur)
		  {
		  ampstim=AMPTETSTIM;
		  }
		if ((time-stime) > (0.0+isi) && (time-stime) < (0.0+isi+cadur))
		  {
		  casyn=AMPTETCA;
		  }
		if ((time-stime) > (0.0+isi) && (time-stime) < (0.0+isi+dcamp))
		  {
		  camp=AMPTETCMP;
		  }
		if ((time-stime) > (0.0+isi) && (time-stime) < (0.0+isi+rasdur))
		  {
		  ampstim=AMPTETSTIM;
		  }
		if ((time-stime) > (0.0+2.0*isi) && (time-stime) < (0.0+2.0*isi+cadur))
		  {
		  casyn=AMPTETCA;
		  }
		if ((time-stime) > (0.0+2.0*isi) && (time-stime) < (0.0+2.0*isi+dcamp))
		  {
		  camp=AMPTETCMP;
		  }
		if ((time-stime) > (0.0+2.0*isi) && (time-stime) < (0.0+2.0*isi+rasdur))
		  {
		  ampstim=AMPTETSTIM;
		  }

// FOLLOWING PARAMETERS CAN BE USED TO SIMULATE KINASE INHIBITOR APPLICATIONS.

		inherk=1.0;
		inhck2=1.0;
		inhgp=1.0;
		inhpka=1.0;
		inhmkk=1.0;
		inhpkm=1.0;

/*		tref = time-inhtime; // this code can be used for simulations of Fig. 4.

		if (tref > 0.0 && tref < inhdur)
		  {
		  inhpkm=0.1;
		  }
*/
		powca=casyn*casyn*casyn*casyn;
		powkc=Kck2*Kck2*Kck2*Kck2;
		termcamp=camp*camp/(camp*camp+Kcamp*Kcamp);

		rafp=raftot-raf;
		mkkp=mkktot-mkk-mkkpp;
		erkp=erktot-erk-erkpp;

dv1dt = kfck2*(powca/(powca+powkc)) - ck2act/tauck2;

dv2dt = (termcamp - pkaact)/taupka;

dv3dt = -(ampstim+kfbasraf)*raf+kbraf*rafp;

dv4dt = -inhmkk*kfmkk*rafp*mkk/(mkk+Kmkk)+kbmkk*mkkp/(mkkp+Kmkk);

dv5dt = inhmkk*kfmkk*rafp*mkkp/(mkkp+Kmkk)-kbmkk*mkkpp/(mkkpp+Kmkk);

dv6dt = -kferk*mkkpp*erk/(erk+Kmk)	+kberk*erkp/(erkp+Kmk);

dv7dt = kferk*mkkpp*erkp/(erkp+Kmk)-kberk*erkpp/(erkpp+Kmk);

		erkact = erkpp;

dv8dt = inhck2*kphos1*ck2act*(1.0-tagp1) - kdeph1*tagp1;

dv9dt = inhpka*kphos2*pkaact*(1.0-tagp2) - kdeph2*tagp2;

dv10dt = inherk*kphos3*erkact*(1.0-tagp3) - kdeph3*tagp3;

		tagsyn = tagp1*tagp2*tagp3;

dv11dt = inhgp*vsyngp - prp/taugp;

dv12dt = -kprew*tagsyn*prp*(prew/(prew+Kpr2)) + vprew - prew/tauprew;

dv13dt = kltp*tagsyn*prp*(prew/(prew+Kpr2))*inhpkm*pkms + vsynbas - wsyn/tausyn;

dv14dt = kphos4*inhck2*ck2act*(1.0-pck2)-kdeph4*pck2;

dv15dt = kphos5*inherk*erkact*(1.0-perk)-kdeph5*perk;

dv16dt = ktranspkm1*pck2*perk + vbaspkm - kdegpkm*pkms;

		values[1]+=dt*dv1dt;
		values[2]+=dt*dv2dt;
		values[3]+=dt*dv3dt;
		values[4]+=dt*dv4dt;
		values[5]+=dt*dv5dt;
		values[6]+=dt*dv6dt;
		values[7]+=dt*dv7dt;
		values[8]+=dt*dv8dt;
		values[9]+=dt*dv9dt;
		values[10]+=dt*dv10dt;
		values[11]+=dt*dv11dt;
		values[12]+=dt*dv12dt;
		values[13]+=dt*dv13dt;
		values[14]+=dt*dv14dt;
		values[15]+=dt*dv15dt;
		values[16]+=dt*dv16dt;

/* Equilibrate basal synaptic weight, prior to stimulation, according to basal kinase activities and other variables. necessary because W is the slowest variable.
For reactivation simulations, omit this prior equilibration because W should fluctuate around basal level before stimulus. Need to initialize it near basal level though. */

/*		if (time < stime)
		  {
		  wsyn = tausyn*(kltp*tagsyn*prp*(prew/(prew+Kpr2))*inhpkm*pkms+vsynbas);
		  values[13] = tausyn*(kltp*tagsyn*prp*(prew/(prew+Kpr2))*inhpkm*pkms+vsynbas);

		  wbas = wsyn;
		  }				*/

// Increment time
                time=time+dt;
                treact=treact+dt;

// END INNER LOOP

                j++;
               } while (j <= delta/dt);

// COMPUTE AND PRINT OUTPUT VARIABLES

            if ((time > recstart) && (time < recend) && (k % recintvl == 0))
              {

/* OUTPUT CONCENTRATION UNITS ARE uM. SCALING FACTORS ARE FOR EASE OF CONCURRENT VISUALIZATION */

/* NOTE – CAMKII is not sampled often enough here to pick up its activation peaks reliably during the reactivation simulations.  A separate simulation that only tracks CaMKII, with higher sampling resolution, was done to better fix its peaks. */

               out1.println(tref + "\t" + 0.12*ck2act);
               out2.println(tref + "\t" + 11.0*erkact);
               out3.println(tref + "\t" + 50.0*pkaact);
               out4.println(tref + "\t" + 0.1*wsyn);
               out5.println(tref + "\t" + 1000.0*tagsyn);
               out6.println(tref + "\t" + 2.0*prew);
               out7.println(tref + "\t" + tagp1);
               out8.println(tref + "\t" + tagp2);
               out9.println(tref + "\t" + tagp3);
               out10.println(tref + "\t" + casyn);
               out11.println(tref + "\t" + 10.0*rafp);
               out12.println(tref + "\t" + pck2);
               out13.println(tref + "\t" + perk);
               out14.println(tref + "\t" + 1.5*pkms);

/* For simulations with inhibitor, use “inhpkm*pkms” instead of just “pkms” in last print line, to print out pkm timecourse with inhibition */

/*  These are scaling factors for simple LTP simulation, no feedback.
               out1.println(tref + "\t" + 0.12*ck2act);
               out2.println(tref + "\t" + 7.0*erkact);
               out3.println(tref + "\t" + 50.0*pkaact);
               out4.println(tref + "\t" + 0.1*wsyn);
               out5.println(tref + "\t" + 700.0*tagsyn);
               out6.println(tref + "\t" + 2.0*prew);
               out7.println(tref + "\t" + tagp1);
               out8.println(tref + "\t" + tagp2);
               out9.println(tref + "\t" + tagp3);
               out10.println(tref + "\t" + casyn);
               out11.println(tref + "\t" + 10.0*rafp);
               out12.println(tref + "\t" + pck2);
               out13.println(tref + "\t" + perk);
               out14.println(tref + "\t" + 2.0*pkms);

For bistable CaMKII simulation, the scale factors are the same EXCEPT FOR PKM. There, we drop from 2.0 down to 0.3 for display.

For bistable PKM simulation, the scale factors are again the same except, we move the PKM one back up to 0.6.

For reactivation feed simulation, these are the scale factors,

               out1.println(tref + "\t" + 0.12*ck2act);
               out2.println(tref + "\t" + 3.5*erkact);
               out3.println(tref + "\t" + 50.0*pkaact);
               out4.println(tref + "\t" + 0.1*wsyn);
               out5.println(tref + "\t" + 240.0*tagsyn);
               out6.println(tref + "\t" + 2.0*prew);
               out7.println(tref + "\t" + tagp1);
               out8.println(tref + "\t" + tagp2);
               out9.println(tref + "\t" + tagp3);
               out10.println(tref + "\t" + casyn);
               out11.println(tref + "\t" + 10.0*rafp);
               out12.println(tref + "\t" + pck2);
               out13.println(tref + "\t" + perk);
               out14.println(tref + "\t" + 0.6*pkms);

*/

              }

            k++;
           } while (k <= recend/delta);

// CLOSE OUTPUT FILES.

        out1.close();
        out2.close();
        out3.close();
        out4.close();
        out5.close();
        out6.close();
        out7.close();
        out8.close();
        out9.close();
        out10.close();
        out11.close();
        out12.close();
        out13.close();
        out14.close();

       }
}
```
