## supplemental java program 1 for "Comparing Theories for the Maintenance of Late LTP and Long-Term Memory: Computational Analysis of the Roles of Kinase Feedback Pathways and Synaptic Reactivation"

```
import java.io.*;
import java.util.Random;

public class fig2ab3a

/* To compile and run these supplemental java programs, install Java Development Kit (JDK) from Oracle, or a similar compiler. Rename this file as fig2ab3a.java. With JDK, type “javac fig2ab3a.java” to compile and “java fig2ab3a” to run. */

 {  public static void main (String args[]) throws IOException {

    PrintWriter out1 = new PrintWriter(new FileWriter("ck2act.txt"));
    PrintWriter out2 = new PrintWriter(new FileWriter("erkact.txt"));
    PrintWriter out3 = new PrintWriter(new FileWriter("pkaact.txt"));
    PrintWriter out4 = new PrintWriter(new FileWriter("wsyn.txt"));
    PrintWriter out5 = new PrintWriter(new FileWriter("tagsyn.txt"));
    PrintWriter out6 = new PrintWriter(new FileWriter("prew.txt"));
    PrintWriter out7 = new PrintWriter(new FileWriter("tagp1.txt"));
    PrintWriter out8 = new PrintWriter(new FileWriter("tagp2.txt"));
    PrintWriter out9 = new PrintWriter(new FileWriter("tagp3.txt"));
    PrintWriter out10 = new PrintWriter(new FileWriter("casyn.txt"));
    PrintWriter out11 = new PrintWriter(new FileWriter("rafp.txt"));
    PrintWriter out12 = new PrintWriter(new FileWriter("pck2.txt"));
    PrintWriter out13 = new PrintWriter(new FileWriter("perk.txt"));
    PrintWriter out14 = new PrintWriter(new FileWriter("pkms.txt"));

    double dt=0.000025; // high-res timestep (min)
    double delta=0.01; // low-res timestep (min)


/* The following timing parameters can be adjusted as needed to reproduce simulations in Figs. 2A, 2B, or 3A */

double recstart=3000.0; // Time to start writing data

    double stime=recstart+179.999; // Time of stimulus onset

    double isi=5.0;      // Spacing between tetanic stimuli

    double recend=stime+899.999;

    int recintvl=20; // (delta) Intervals to record

	double tref;

//PROGRAM VARIABLES

// Activities of kinases other than MAPK cascades
	double ck2act;
	double camp;
	double pkaact;

// MAPK CASCADE
	double raf;
	double rafp;
	double mkk;
	double mkkp;
	double mkkpp;
	double erk;
	double erkp;
	double erkpp;
	double erkact;

// SYNAPTIC VARIABLES
	double tagp1;
	double tagp2;
	double tagp3;
	double tagsyn;
	double prew;
	double wsyn;
	double wbas=0.0;

// SIMPLE PLASTICITY RELATED PROTEIN
	double prp;

// pkm level, and phosphorylation sites on TFs for CaMKII and ERK

	double pck2;
	double perk;
	double pkms;
	double pkmfterm;  // term for PKM autoactivation

// TIME DERIVATIVES

    	double dv1dt;
    	double dv2dt;
    	double dv3dt;
    	double dv4dt;
    	double dv5dt;
    	double dv6dt;
    	double dv7dt;
    	double dv8dt;
    	double dv9dt;
    	double dv10dt;
    	double dv11dt;
    	double dv12dt;
    	double dv13dt;
    	double dv14dt;
	double dv15dt;
	double dv16dt;
	double dv17dt;
	double dv18dt;
	double dv19dt;
	double dv20dt;

// Cytoplasmic and nuclear Ca concentrations, camp function
	double casyn;
	double powca;
	double powkc;
	double termcamp;

// WORK IN CONCENTRATION UNITS OF uM, TIME UNITS OF MIN

// MODEL PARAMETERS FOLLOW:

// BASAL PKA AND CA, STIMULUS AMPLITUDES. CA2 refers to nuclear Ca.
// Stim amps also drive PKA elevations.

	double Cabas=0.04;
	double campbas=0.06;
	double AMPTETCA=0.8;
	double cadur=0.05;

	double AMPTETCMP=0.25;
	double dcamp=1.0; // duration of tetanic cAMP elevation. 1.0 min standard RTS, rolipram cures RTS-basal by raising dcamp to 2.0.

	double AMPTETSTIM=0.13;

	double rasdur=1.0; // 1.0 standard

	double ampstim; // set later to Raf activation stimulus amplitude

// RATE CONSTANTS AND MICH CONSTANT FOR CAMKIV AND CAMKII ACTIVATION
	double kfck2=180.0; // 180 standard for most sims

	double tauck2=1.0;
	double Kck2=0.7;

//CONSERVED ENZYME AMOUNTS
	double raftot=0.25;
	double mkktot=0.25;
	double erktot=0.25;

// MAPK rate and Michaelis constants
	double kbraf=0.12;
	double kfmkk=0.6;
	double kbmkk=0.025;
	double Kmkk=0.25;
	double kferk=0.52;
	double kberk=0.025;
	double Kmk=0.25;
	double kfbasraf=0.0075; /* 0.0075 standard for most sims */

// Inhibition factors for enzymes and for plasticity related protein gene

	double inherk;
	double inhmkk;
	double inhck2;
	double inhgp;
	double inhpka;
	double inhpkm;

// Rate constants and other parameters
// describing PKA activation and synaptic tagging

	double Kcamp=1.0;

	double kphos1=0.15;
	double kdeph1=0.008;
	double kphos2=0.8;
	double kdeph2=0.2;
	double kphos3=0.06;
	double kdeph3=0.05;
	double taupka=15.0;

// PKM parameters

	double kphos4=0.1;
	double kdeph4=0.1;
	double kphos5=2.0;
	double kdeph5=0.1;
	double kdegpkm=0.02;

	double ktranspkm1=0.2;

	double Kpkm=0.75;
	double vbaspkm=0.0015;

	double epsilpkm=0.0; /* coupling constant to tune the PKM positive feedback strength, I find that 0.028 is the number I want for bistability here (the equations are simplified and modified from the 2012 paper, so easier to get bistability) */

	double epsilck2=0.0; // coupling constant to tune strength of CaMKII autoactivation positive feedback loop.  1.0 gives strong bistability.

	double kfeedck2=4.0; // two more parameters for quadratic CaMKII feedback loop
	double Khck2=1.0;

// Maximal and basal transcription rates for transcription of plasticity related protein (could also be interpreted as translation), and for decay of PRP.

// Note that inhibition factor for protein synthesis or transcription is given a value here

	double vsyngp=0.01;

	inhgp=1.0;

	double taugp=100.0;

// Rate constants and other parameters for synaptic weight changes

/* kltp=500, Kpr2=0.2, vprew=0.0035, for single decaying tetanic LTP, no feedback. */

/* for CaMKII feedback simulation, same values except kltp is adjusted down to 70. */

/* For PKM only feedback simulation, need kltp back up. Is up to 300. */

/* kltp = 100 is standard value for generating bistability in W with the upper state stabilized by synaptic reactivation. */

	double kltp=500.0;

	double vsynbas = 0.01;

	double tausyn=300.0; // 300 is standard value.

	double kprew=6.0;

	double Kpr2=0.2; // Dissociation type constant by which a decrease in PREW protein suppresses rate of increase of W.

	double vprew=0.0035;

	double tauprew=100.0;


// TIME STUFF, COUNTERS, VALUE ARRAY

    double time=0.0; // time (minutes)

    int i, k, j, l, m, iout, ibig; // counters

    double[] values=new double[20];


// VARIABLE INITIALIZATION. Use nonzero initial values
// to avoid extremely small numbers in output files.

/* for slower variables, initial conditions are set reasonably close to equilibrium basal values, for fast variables 0.001 is fine */

	values[1]=0.001;
	values[2]=0.001;
	values[3]=0.5*raftot;
	values[4]=0.3*mkktot;
	values[5]=0.4*mkktot;
	values[6]=0.3*erktot;
	values[7]=0.4*erktot;
	values[8]= 0.001;
	values[9]= 0.001;
	values[10]= 0.001;
	values[11]=inhgp*vsyngp*taugp;
	values[12]=vprew*tauprew;
	values[13]=vsynbas*tausyn;
	values[14]= 0.001;
	values[15]= 0.001;
	values[16]= 0.001;

// MAIN LOOP (LARGER TIMESTEP, IF TIME COURSE DATA IS OUTPUTTED THIS LARGER TIMESTEP IS USED)

        k=1;
        do {

// INNER SIMULATION LOOP (SMALLER TIMESTEP DT)

            j=1;
            do {

	/* code for inhibition simulation	tref=time-inhtime; */

		tref=time-stime;
		casyn=Cabas;
		ampstim=0.0;
		camp=campbas;

		ck2act=values[1];
		pkaact=values[2];

		raf=values[3];
		mkk=values[4];
		mkkpp=values[5];
		erk=values[6];
		erkpp=values[7];

		tagp1=values[8];
		tagp2=values[9];
		tagp3=values[10];

		prp=values[11];

		prew=values[12];
		wsyn=values[13];

		pck2=values[14];
		perk=values[15];
		pkms=values[16];


// SIMPLE SQUARE WAVES FOR CYTOPLASMIC CA
// DURING THE TETANIC PROTOCOL.

/* note that several LTP stimulus protocols, including chem-LTP and theta-LTP, can be simulated here. See Smolen et al 2006 for these other protocols */

// Tetanic protocol

		if ((time-stime) > 0.0 && (time-stime) < cadur)
		  {
		  casyn=AMPTETCA;
		  }
		if ((time-stime) > 0.0 && (time-stime) < dcamp)
		  {
		  camp=AMPTETCMP;
		  }
		if ((time-stime) > 0.0 && (time-stime) < rasdur)
		  {
		  ampstim=AMPTETSTIM;
		  }
		if ((time-stime) > (0.0+isi) && (time-stime) < (0.0+isi+cadur))
		  {
		  casyn=AMPTETCA;
		  }
		if ((time-stime) > (0.0+isi) && (time-stime) < (0.0+isi+dcamp))
		  {
		  camp=AMPTETCMP;
		  }
		if ((time-stime) > (0.0+isi) && (time-stime) < (0.0+isi+rasdur))
		  {
		  ampstim=AMPTETSTIM;
		  }
		if ((time-stime) > (0.0+2.0*isi) && (time-stime) < (0.0+2.0*isi+cadur))
		  {
		  casyn=AMPTETCA;
		  }
		if ((time-stime) > (0.0+2.0*isi) && (time-stime) < (0.0+2.0*isi+dcamp))
		  {
		  camp=AMPTETCMP;
		  }
		if ((time-stime) > (0.0+2.0*isi) && (time-stime) < (0.0+2.0*isi+rasdur))
		  {
		  ampstim=AMPTETSTIM;
		  }

// FOLLOWING PARAMETERS CAN BE USED TO SIMULATE KINASE INHIBITOR APPLICATIONS.

		inherk=1.0;
		inhck2=1.0;
		inhgp=1.0;
		inhpka=1.0;
		inhmkk=1.0;
		inhpkm=1.0;

/*		tref = time-inhtime; // this code can be used for simulations of Fig. 4.

		if (tref > 0.0 && tref < inhdur)
		  {
		  inhpkm=0.1;
		  }
*/
		powca=casyn*casyn*casyn*casyn;
		powkc=Kck2*Kck2*Kck2*Kck2;
		termcamp=camp*camp/(camp*camp+Kcamp*Kcamp);

		rafp=raftot-raf;
		mkkp=mkktot-mkk-mkkpp;
		erkp=erktot-erk-erkpp;

dv1dt = kfck2*(powca/(powca+powkc)) - ck2act/tauck2 + epsilck2*(kfeedck2/tauck2*ck2act*ck2act/(Khck2*Khck2+ck2act*ck2act));

dv2dt = (termcamp - pkaact)/taupka;

dv3dt = -(ampstim+kfbasraf)*raf+kbraf*rafp;

dv4dt = -inhmkk*kfmkk*rafp*mkk/(mkk+Kmkk)+kbmkk*mkkp/(mkkp+Kmkk);

dv5dt = inhmkk*kfmkk*rafp*mkkp/(mkkp+Kmkk)-kbmkk*mkkpp/(mkkpp+Kmkk);

dv6dt = -kferk*mkkpp*erk/(erk+Kmk)	+kberk*erkp/(erkp+Kmk);

dv7dt = kferk*mkkpp*erkp/(erkp+Kmk)-kberk*erkpp/(erkpp+Kmk);

		erkact = erkpp;

dv8dt = inhck2*kphos1*ck2act*(1.0-tagp1) - kdeph1*tagp1;

dv9dt = inhpka*kphos2*pkaact*(1.0-tagp2) - kdeph2*tagp2;

dv10dt = inherk*kphos3*erkact*(1.0-tagp3) - kdeph3*tagp3;

		tagsyn = tagp1*tagp2*tagp3;

dv11dt = inhgp*vsyngp - prp/taugp;

dv12dt = -kprew*tagsyn*prp*(prew/(prew+Kpr2)) + vprew - prew/tauprew;

dv13dt = kltp*tagsyn*prp*(prew/(prew+Kpr2))*inhpkm*pkms + vsynbas - wsyn/tausyn;

dv14dt = kphos4*inhck2*ck2act*(1.0-pck2)-kdeph4*pck2;

dv15dt = kphos5*inherk*erkact*(1.0-perk)-kdeph5*perk;

pkmfterm = pkms*pkms/(pkms*pkms+Kpkm*Kpkm);

dv16dt = ktranspkm1*pck2*perk + vbaspkm + epsilpkm*pkmfterm - kdegpkm*pkms;

		values[1]+=dt*dv1dt;
		values[2]+=dt*dv2dt;
		values[3]+=dt*dv3dt;
		values[4]+=dt*dv4dt;
		values[5]+=dt*dv5dt;
		values[6]+=dt*dv6dt;
		values[7]+=dt*dv7dt;
		values[8]+=dt*dv8dt;
		values[9]+=dt*dv9dt;
		values[10]+=dt*dv10dt;
		values[11]+=dt*dv11dt;
		values[12]+=dt*dv12dt;
		values[13]+=dt*dv13dt;
		values[14]+=dt*dv14dt;
		values[15]+=dt*dv15dt;
		values[16]+=dt*dv16dt;

/* Equilibrate basal synaptic weight, prior to stimulation, according to basal kinase activities and other variables. necessary because W is the slowest variable */

		if (time < stime)
		  {
		  wsyn = tausyn*(kltp*tagsyn*prp*(prew/(prew+Kpr2))*inhpkm*pkms+vsynbas);
		  values[13] = tausyn*(kltp*tagsyn*prp*(prew/(prew+Kpr2))*inhpkm*pkms+vsynbas);

		  wbas = wsyn;
		  }

// Increment time
                time=time+dt;

// END INNER LOOP

                j++;
               } while (j <= delta/dt);

// COMPUTE AND PRINT OUTPUT VARIABLES

            if ((time > recstart) && (time < recend) && (k % recintvl == 0))
              {

/* OUTPUT CONCENTRATION UNITS ARE uM. SCALING FACTORS ARE FOR EASE OF CONCURRENT VISUALIZATION */

               out1.println(tref + "\t" + 0.12*ck2act);
               out2.println(tref + "\t" + 7.0*erkact);
               out3.println(tref + "\t" + 50.0*pkaact);
               out4.println(tref + "\t" + 0.1*wsyn);
               out5.println(tref + "\t" + 700.0*tagsyn);
               out6.println(tref + "\t" + 2.0*prew);
               out7.println(tref + "\t" + tagp1);
               out8.println(tref + "\t" + tagp2);
               out9.println(tref + "\t" + tagp3);
               out10.println(tref + "\t" + casyn);
               out11.println(tref + "\t" + 10.0*rafp);
               out12.println(tref + "\t" + pck2);
               out13.println(tref + "\t" + perk);
               out14.println(tref + "\t" + 2.0*pkms);

/* For simulations with inhibitor, use “inhpkm*pkms” instead of just “pkms” in last print line, to print out pkm timecourse with inhibition */

/*  These are scaling factors for simple LTP simulation, no feedback.
               out1.println(tref + "\t" + 0.12*ck2act);
               out2.println(tref + "\t" + 7.0*erkact);
               out3.println(tref + "\t" + 50.0*pkaact);
               out4.println(tref + "\t" + 0.1*wsyn);
               out5.println(tref + "\t" + 700.0*tagsyn);
               out6.println(tref + "\t" + 2.0*prew);
               out7.println(tref + "\t" + tagp1);
               out8.println(tref + "\t" + tagp2);
               out9.println(tref + "\t" + tagp3);
               out10.println(tref + "\t" + casyn);
               out11.println(tref + "\t" + 10.0*rafp);
               out12.println(tref + "\t" + pck2);
               out13.println(tref + "\t" + perk);
               out14.println(tref + "\t" + 2.0*pkms);

For bistable CaMKII simulation, the scale factors are the same EXCEPT FOR PKM. There, we drop from 2.0 down to 0.3 for display.

For bistable PKM simulation, the scale factors are again the same except, we move the PKM one back up to 0.6.

*/

              }

            k++;
           } while (k <= recend/delta);

// CLOSE OUTPUT FILES.

        out1.close();
        out2.close();
        out3.close();
        out4.close();
        out5.close();
        out6.close();
        out7.close();
        out8.close();
        out9.close();
        out10.close();
        out11.close();
        out12.close();
        out13.close();
        out14.close();

       }
}
```
